# Testing the Drosophila *maternal haploid* gene for functional divergence and a role in hybrid incompatibility

**DOI:** 10.1101/2022.03.23.485453

**Authors:** Dean M Castillo, Connor M Kean, Benjamin McCormick, Sahana Natesan, Daniel A Barbash

**Author notes:** Correspondence to: Dean Castillo or Daniel Barbash.

## Abstract

Crosses between *D. simulans* females and *D. melanogaster* males produce viable F1 sons and poorly viable F1 daughters. Unlike most hybrid incompatibilities, this hybrid incompatibility violates Haldane’s rule, the observation that incompatibilities preferentially affect the heterogametic sex. Further, it is important to understand fully the genetic basis of this incompatibility because the causal allele in *D. melanogaster* is a large species-specific block of complex satellite DNA on its X chromosome known as the 359-bp satellite, rather than a protein-coding locus. The causal allele(s) in *D. simulans* are unknown but likely involve maternally expressed genes or factors since the F1 females die during early embryogenesis. The *maternal haploid* (*mh*) gene is an intriguing candidate because it is expressed maternally and its protein product localizes to the 359-bp repeat. We found that this gene has diverged extensively between *D. melanogaster* and *D. simulans*. This observation led to the hypothesis that *D. melanogaster mh* may have co-evolved with the 359-bp repeat, and that hybrid incompatibility thus results from the absence of a co-evolved *mh* allele in *D. simulans*. We tested for functional divergence of *mh* by creating matched transformants of *D. melanogaster* and *D. simulans* orthologs in both *D. melanogaster* and *D. simulans* strains. Surprisingly, we find that *D. simulans mh* fully complements the female sterile phenotype of *D. melanogaster mh* mutations. Contrary to our hypothesis, we find no evidence that adding a *D. melanogaster mh* gene to *D. simulans* increases hybrid viability.

## Introduction

The evolution of reproductive isolation via hybrid incompatibilities can be complex, with multiple incompatibilities contributing to isolation within a single species pair. Two genetically distinct lethal hybrid incompatibilities exist between the sister species *D. melanogaster* and *D. simulans* (Sawamura *et al*. 1993b; Barbash 2010). When *D. melanogaster* females are crossed to *D. simulans* males, the F1 hybrid sons are invariably lethal, while the F1 daughters are generally fully viable, at least at lower temperatures (~<25°) (Sturtevant 1920; Barbash *et al*. 2000). Importantly, this pattern of lethality is not sex-specific but rather caused by the presence of the *D. melanogaster* X chromosome. Experiments that can detect products of non-disjunction or use attached-X chromosomes demonstrate that daughters inheriting both X chromosomes from their *D. melanogaster* mother are lethal while sons inheriting their X from their *D. simulans* father are viable (Barbash 2010). The lethality is caused by an incompatibility between the *D. melanogaster* allele of the gene *Hmr* on the *D. melanogaster* X, interacting with the *D. simulans* alleles of the autosomal genes *Lhr* and *GFZF* (Watanabe 1979; Hutter and Ashburner 1987; Brideau *et al*. 2006; Phadnis *et al*. 2015).

In contrast, the reciprocal cross of *D. simulans* females to *D. melanogaster* males produces viable F1 sons and poorly viable F1 daughters that die as early embryos (Sturtevant 1920). While the *D. melanogaster* X is again implicated in causing this embryonic lethality, Sawamura et al. (1993c) showed that *Hmr* is not responsible for this F1 lethality. Instead, the lethal effect of the *D. melanogaster* X maps to the pericentromeric heterochromatin in a region called *Zhr* (Sawamura and Yamamoto 1993). The lethal effect of *Zhr*^+^ appears to be caused by missegregation during early embryogenesis of a multi-million base pair block of complex satellite DNA sequences (Ferree and Barbash 2009). These satellite sequences are known alternatively as the 359-bp or the 1.688 g/cm^3^ satellites (Sawamura and Yamamoto 1993; Ferree and Barbash 2009) (Lohe and Brutlag 1986). While *D. simulans* contains some dispersed 359-bp repeats, it does not have the large X-linked block found in *D. melanogaster* (Lohe *et al*. 1993; Sproul *et al*. 2020; de Lima *et al*. 2020). This extensive difference in abundance of the 359-bp satellite between the species suggests that *D. melanogaster* may contain allele(s) that have coevolved with the X-linked 359-bp satellite block to help promote its proper mitotic segregation.

This logic further implies that the lack of the X-linked 359-bp satellite block in *D. simulans* would cause it to be unable to regulate this satellite block when inherited from a *D. melanogaster* parent, leading to hybrid incompatibility. Because the F1 hybrid female lethality occurs during early embryonic development, the allele(s) hypothesized to be missing from *D. simulans* are likely to be maternally expressed (Sturtevant 1920; Ferree and Barbash 2009). The penetrance of F1 hybrid female lethality is highly variable across strains, which has complicated efforts to identify the genes causing this lethality. Sawamura et al. identified a strain of *D. simulans* called *maternal hybrid rescue* (*mhr*) that produces high viability of F1 hybrid daughters and mapped the effect to the second chromosome (Sawamura *et al*. 1993a). Orr also implicated the *D. simulans* second chromosome in contributing to hybrid lethality in this cross (Orr 1996). Another study directly tested the satellite-binding protein D1 as a candidate but found no evidence for a role in the incompatibility (Ferree and Barbash 2009). Further attempts to identify the genetic basis of the *D. simulans* maternal effect on hybrid viability led to the plausible conclusion that it is a polygenic effect (Gérard and Presgraves 2012). Others have suggested that the incompatibility may be caused by the absence in *D. simulans* of maternally deposited small RNAs homologous to the 359-bp repeat, rather than by protein-coding genes (Ferree and Barbash 2007). Several studies have shown that such RNAs are produced by heterochromatic satellites including the X-linked 359-bp satellite block (Usakin *et al*. 2007; Wei *et al*. 2021), though their potential role in hybrid lethality remains untested.

The X-linked gene *maternal haploid* (*mh*) is an intriguing candidate for contributing to this interspecific incompatibility. Mutations in *mh* were first identified based on its female sterility phenotype (Gans *et al*. 1975). Embryos from *mh* mutant mothers typically arrest within the first few nuclear cycles with condensation defects specific to the paternally inherited chromosomes, with a minority reaching late embryogenesis as lethal gynogenetic haploids (Zalokar *et al*. 1975; Loppin *et al*. 2001). The *mh* gene encodes a predicted metalloprotease, homologs of which are involved in DNA damage repair (Delabaere *et al*. 2014; Tang *et al*. 2017). Most relevant to this study, the *mh* protein localizes to the 359-bp satellite during embryogenesis and the satellite shows aberrant segregation in the embryonic progeny of *mh* mutant mothers (Tang *et al*. 2017). These findings, as well as the evolutionary patterns described below, motivated us to test for functional divergence of *mh* orthologs between *D. melanogaster* and *D. simulans* and for possible effects on hybrid lethality.

In particular, we tested the hypothesis that hybrid female embryonic lethality results from the inability of the maternally expressed *D. simulans* Mh protein to properly interact with the paternally inherited X-linked *D. melanogaster* 359-bp satellite block. Under this hypothesis, we predict that adding *mel-mh* to *D. simulans* mothers would suppress hybrid lethality.

## Materials and Methods

### Nomenclature

We use the abbreviations *mel-mh* and *sim-mh* to refer to the *mh* ortholog in *D. melanogaster* and *D. simulans*, respectively. We use *phi{mel-mh-Gfp}* and *phi{sim-mh-Gfp}* to designate PhiC31-mediated transgenes containing Gfp fusions of *mel-mh* and *sim-mh*, respectively. As described in the Results, *D. simulans* has a tandem duplication of *mh*. The *mh-p* copy is more similar than the *mh-d* copy in sequence and structure to *D. melanogaster mh*. We therefore consider *sim-mh-p* to be the ortholog of *D. melanogaster mh*, and for simplicity refer to it as *sim-mh*.

### Drosophila stocks

*w mh^6^* / *FM7a, P{sChRFP}1* and *w, mh^31^/FM7, GFP^+^* stocks were kindly provided by Xiaona Tang and Yikang Rong. *D. simulans* stocks containing attP landing sites were kindly provided by David Stern (Stern *et al*. 2017).

### Sequence analysis

The sequences for both duplicates of *mh* from the *D. simulans* genome were taken from the genome sequence and aligned with the *D. melanogaster* ortholog to determine the consensus coding sequence (see Results). We then calculated D_N_/D_S_ between the *D. melanogaster mh* and both *D. simulans* orthologs using MEGA 11 (Tamura *et al*. 2021). We compared these ratios to a genome-wide sample of D_N_/D_S_ ratios from a previously published study to determine their percentile and relative rate of evolutionary change (Stanley and Kulathinal 2016). These data all consist of two-by-two (i.e. pairwise) sequence comparisons between *D. melanogaster* and *D. simulans* orthologs.

### *mh* transgene constructs

The *mel-mh-gfp* transgene was kindly provided by Xiaona Tang and Yikang Rong and previously described as *gfp-mh-pTV2gw* (Tang *et al*. 2017). We added an *attB* site to this to create the plasmid *gfp-mh-pTV2gw-attB* by PCR-amplifying using oligos 502/503 from a plasmid with an *attB* site that derived from the *pTA-attB* plasmid (Groth *et al*. 2004). The PCR product was digested using NotI and inserted into *gfp-mh-pTV2gw* at its NotI site. All oligonucleotide sequences are listed in Table S1.

The *w+-attB-sim-mh-eGFP* (p834) construct was designed to be parallel in structure to *gfp-mh-pTV2gw-attB* (Fig 1 and Sup File 1) and was made in the following steps:

1. *w+-attB-sim-mh*. The *D. simulans mh* genomic region covering the coding region and ~3000 bp upstream was PCR amplified from the strain *w^501^* using oligos 2099/2100 and cloned into pCR-Blunt II-TOPO (Invitrogen). The sequence of *sim-mh* was confirmed by Sanger sequencing (using oligos 788, 823, 2078, 2079, 2080, 2081, 2082, 2083, 2084 and 2101). The insert was then released by XbaI digestion and ligated into the XbaI site of the plasmid *w^+^-attB*, a gift from Jeff Sekelsky (Addgene plasmid # 30326; http://n2t.net/addgene:30326; RRID:Addgene_30326). The resulting clone *w^+^-attB-sim-mh* was confirmed by checking the pattern of restriction enzyme digestion.
2. *pCR-Blunt II-TOPO_sim-mh(partial)-eGFP*. The coding sequence of eGFP was inserted into the coding sequence of *sim-mh* immediately after the start codon by Gibson assembly of Dsmh1, eGFP and Dsmh2. Dsmh1 and Dsmh2 were PCR amplified from *D. simulans* (strain *w^501^*) using oligo pairs 2016/2017and 2020/2021, respectively, while eGFP was amplified using oligos 2018/2019 from pEGFP-attB (Drosophila Genomics Resource Center). A fusion product of Dsmh1-eGFP -Dsmh2 was PCR-amplified from the Gibson Assembly reaction mixture, using oligos 2080/2082 and cloned into pCR-Blunt II-TOPO. The sequence of the insert, i.e. the fusion of eGFP CDS and Dsmh (partial), was checked by Sanger sequencing using oligos 498, 788, 823, 2081.
3. *w+-attB-Dsmh-eGFP*. The Dsmh(partial)_eGFP fragment was released from the pCR-Blunt II TOPO vector and ligated into w+-attB-sim-mh using double digestion (BsiWI-BstEII). The final construct w+attB-sim_mh-eGFP was confirmed by checking the restriction pattern of the construct.

**Fig 1.**
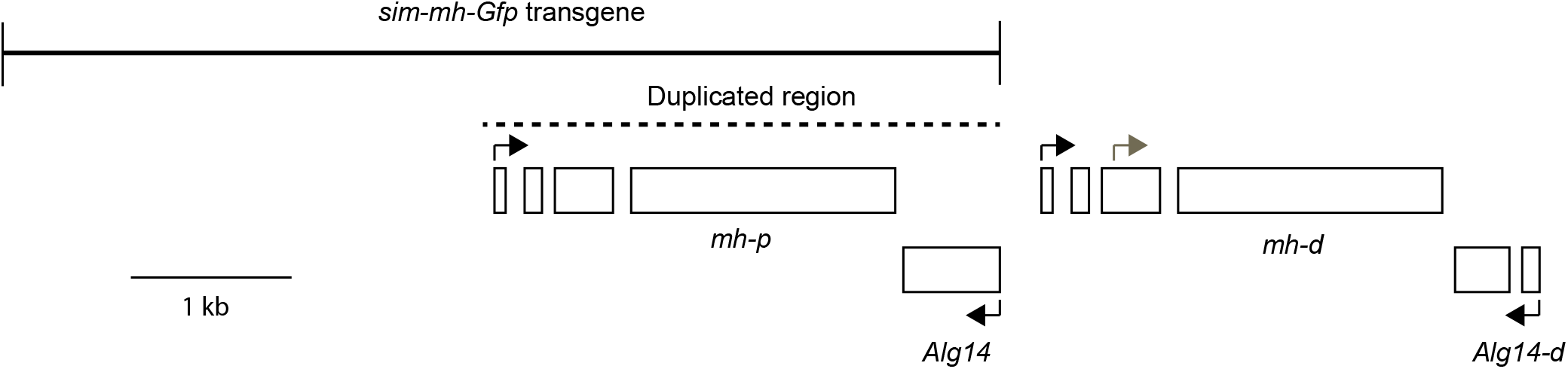
Map of *mh* region in *D. simulans* demonstrating the duplication event. The region in *D. melanogaster* has the same structure as the *mh-p* and *Alg14* region. The dashed line shows the region duplicated to form *mh-d* and *Alg14-d*. Black arrows show translation start sites; the gray arrow in *mh-d* shows the alternative translation start site proposed by (Chakraborty *et al*. 2021). The two boxes shown for *Alg14-d* represent a frameshift relative to *Alg14*. A full annotation of this region is shown in Supp File 1.

### *mh* transgenic lines

*D. melanogaster* transgenic lines were made by phiC31-mediated integration into the strain *y w P{nos-phiC31\int.NLS}X; P{CaryP}attP40*. *D. simulans* transformant lines were made by phiC31-mediated integration into the strains *y w; pBac{3XP3::EYFP,attP}1048-2R* and *y w; pBac{3XP3::EYFP,attP}1029-3R* (Stern *et al*. 2017). Microinjections were done by Rainbow Transgenic Flies, Inc.

### Fertility tests

The *mel-mh-Gfp* and *sim-mh-Gfp* transgenes transformed into *D. melanogaster* were compared for their ability to complement the female sterility of *mh* null mutations. A smallscale pilot experiment was initially done at room temperature by crossing single virgin *w mh^6^ / w mh^31^; {mh-Gfp, w^+^}attP40/+* females to two DGRP-882 (wild type) males, where *mh-Gfp* represents either of the two transgenes. Two sets were done, the first with females aged 3-4 days-old before mating and the second aged 9-10 days. Vials were cleared after 5 days; if either the female or both males were dead, then the vial was discarded

To generate F1 females to assay in a large-scale experiment, a parental cross was set up of *w mh^6^; {mh-Gfp, w^+^}attP40* females and *mh^31^/Y* males, where *mh-Gfp* represents either of the two transgenes. Virgin F1 daughters of genotype *w mh^6^ / w mh^31^; {mh-Gfp, w^+^}attP40/+* were collected and aged for 3-5 days, followed by test crosses containing one virgin female and two Canton-S (wild type) males. After 4-5 days, parents were flipped to new vials, and flipped again after another 4-5 days. At each flip, if either the female or both males were dead, then the vial was discarded. Otherwise, vials were kept for 16 (27°) or 18 (25°) days and all progeny counted. Three flips were performed at 27° and four at 25°; however very few parents survived until the fourth flip at 25° and thus we only report the first three flips in Fig 3B. The overall time period across all three flips was 12 days at 25° and 12-13 days at 27°. Progeny per day are reported in order to normalize between flips that were 4 or 5 days.

### Hybrid viability tests

The *mel-mh-Gfp* and *sim-mh-Gfp* transgenes transformed into *D. simulans* were assessed for their ability to rescue female viability in F1 *D. simulans/D. melanogaster* hybrids as compared to a matched control group lacking the transgenes. To generate F1 hybrids either carrying or not carrying a *mh-Gfp* transgene, for each transgenic *D. simulans* line, the following parental crosses were set; first, *w^501^* virgin females were crossed to *y w/Y; {mh-Gfp, w+}* males, where *mh-Gfp* represents either of the two transgenes. Virgin daughters of the genotype *w^501^ /y w; {mh-Gfp, w+}/+* collected from this cross were subsequently mated to *w^501^ / Y* males. For each set, 30-40 *y? w/w^501^; {mh-Gfp, w+}/+* and *y? w/w^501^* virgin daughters were separately collected, aged 0-1 days, and mated to 40-50 3-5 day old *D. melanogaster* Canton-S virgin males. The crosses were kept at room temperature (19.4-22.0°) and flipped every 2-4 days until they stopped producing progeny; the adult F1 hybrids were scored for sex.

### Western blots

Young female virgin flies were fed yeast paste 2-3 days prior to dissection. Ovaries were dissected in 0.7% NaCl with 2 pairs of tweezers after flies were anesthetized by carbon dioxide. Ovaries were collected into 1.7 ml microcentrifuge tubes and flash-frozen in liquid nitrogen, then stored at −80°C until all dissections were completed.

To extract proteins, ovaries were ground in SDS sample buffer (62.5 mM Tris pH6.8, 2% SDS, 10% glycerol, 1% ß-mercaptoethanol, 0.05% bromophenol blue), boiled for 3 minutes, and centrifuged at 10,000 rpm for 5 minutes. Proteins were separated via 7.5% SDS-PAGE using a BioRad Mini-PROTEAN Vertical Electro Cell and transferred to PVDF (polyvinylidene difluoride) membrane using BioRad Mini Trans-Blot. The protein standard was Thermo Scientific™ PageRuler™ Prestained Protein Ladder, 10 to 180 kDa.

The protein-bound membrane was blocked in 5% skim milk in TBST (150mM NaCl, 20 mM Tris pH7.5, 0.1% Tween-20) for 1 hour at room temperature, followed by primary antibody incubation at 4°C for 16 hours, and secondary antibody incubation at room temperature for 1 hour. The membrane was washed 3 x 10 minutes in TBST after each antibody incubation. Primary antibodies used were: Anti-GFP Rabbit Polyclonal Antibody (1/5,000, Rockland Immunochemical 600-401-215S) and Monoclonal Mouse Anti-α-Tubulin antibody (1/20,000, Sigma T9026), secondary antibodies: HRP-Goat Anti-Rabbit IgG (H+L) (1/4,000, Jackson 111-035-003) and HRP-Goat Anti-Mouse IgG (H+L) (1/8,000, Jackson 115-035-003). Antibodies were diluted in either 5% skim milk or 5% BSA, in TBST. Signals were detected by applying ECL2 Western Blotting Substrate (Thermo Scientific 80197) to the membrane and exposing it to autoradiography film (VWR 490001-930).

## Results

### *mh* is duplicated in *D. simulans*

When attempting to identify the *mh* ortholog in *D. simulans*, we noticed two different regions with homology to *mel-mh*. The first region was a contig on the X with high similarity but that appeared to be incomplete and possibly contain partial duplications. While pursuing this analysis a PacBio assembly of the *D. simulans* genome reported that *mh* is tandemly duplicated on the X along with the flanking gene *alg14* (Chakraborty *et al*. 2021). Following Chakraborty et al. (2021), we refer to the *D. simulans* duplicates as *mh-p* (mh-proximal) and *mh-d* (mh-distal), though as noted below we suggest that *mh-p* can also be considered the parental copy and *mh-d* the daughter copy.

We confirmed the duplication structure of *D. simulans mh* in an independent assembly of *D. simulans* created using Nanopore sequencing (Miller *et al*. 2018). The Nanopore and PacBio assemblies fully agree in structure, with the exception of a ~25 bp insertion in the PacBio assembly immediately distal to the proximal copy (Supp File 1). The two genome assemblies also contain no differences in the coding sequences of either *mh-p* or *mh-d*.

We used Artemis and the Artemis Comparison Tool to compare and annotate the structures of the *D. melanogaster* and *D. simulans mh* regions (cite) (Sup file 1). We estimate that the duplicated region corresponds to 3160 bp of sequence (relative to *D. melanogaster*). This leaves *D. simulans mh-d* having a duplication of its 3’ region as its 5’ region, which may be responsible for its novel testis-enriched expression pattern reported by Chakraborty et al (2021). There is a 16 bp deletion in the coding region of exon 1 of *mh-d*, relative to *mh-p* and *D. melanogaster mh*. We confirmed that *mh-d* has this deletion in five *D. simulans* strains that were sequenced using Sanger sequencing (Begun *et al*. 2007). This deletion changes the coding potential of exon 1, but it could potentially splice to exon 2 and restore the same reading frame as in *mh-p*. We have annotated *mh-d* to have the same length of its N-terminal region as *mh-p* and *D. melanogaster mh* (Fig. 1; Supp file 1). Charkrobarty et al. (2021) instead annotated the *mh-d* CDS as beginning in what would be its third exon relative to the *mh-p* gene structure, based on the reduced RNA-Seq reads mapping to exons 1 and 2. As there are some RNA-Seq reads that appear to map to the (potential) exons 1 and 2 of *mh-d*, further work is warranted to determine the protein product(s) made by *mh-d*.

Regardless of this uncertainty regarding the N-terminal structure of *mh-d, mh-p* has greater similarity *D. melanogaster mh* in its structure, primary sequence, and expression pattern, suggesting that *mh-p* can be considered to be the parental copy and *mh-d* the daughter copy of the duplication. We thus define *D. simulans mh-p* as the ortholog of *D. melanogaster mh*.

The second region of lesser homology mapped to an intron of the *tkv* gene on chr 2. This region of chr 2 is annotated as the pseudogene CR14033, and was identified as producing siRNAs in testis (Czech *et al*. 2008; Okamura *et al*. 2008) that were noted as homologous to *mh* (Czech *et al*. 2008).

### *mh* coding sequences are rapidly evolving

We calculated pairwise divergence among the *mh* genes in *D. melanogaster* and *D. simulans* (Table 1). The D_N_/D_S_ ratios are relatively high compared to a genome-wide sample of loci, indicating substantial non-synonymous divergence. The comparison between *mel-mh* and *sim-mh-p* had a D_N_/D_S_ ratio of 0.373, placing it in the top 10% of the genome-wide distribution. The comparison between *mel-mh* and *sim-mh-d* had a D_N_/D_S_ ratio of 0.541 which is in the top 5% of the genome distribution. For reference, the hybrid incompatibility loci *Hmr* and *Lhr* are in the top 3% of the distribution. Both D_N_ and D_N_/D_S_ are higher between the paralogs *sim-mh-p* and *sim-mh-d* than between the orthologs *mel-mh* and *sim-mh-p*, though we reiterate that there are uncertainties in the annotation of *sim-mh-d* noted in the previous section.

**Table 1.** Divergence of *mh* genes between *D. melanogaster* and *D. simulans*

| Gene 1 | Gene 2 | D<sub>N</sub> | D<sub>S</sub> | D<sub>N</sub>/D<sub>S</sub> | Percentile |
| --- | --- | --- | --- | --- | --- |
| *mel_mh* | *sim-mh-p* | 0.056 | 0.150 | 0.373 | 91.2 |
| *mel_mh* | *sim-mh-d* | 0.106 | 0.196 | 0.541 | 95.8 |
| *sim-mh-p* | *sim-mh-d* | 0.059 | 0.093 | 0.638 | n/a |

The numbers of non-synonymous and synonymous substitutions per site were estimated using the Nei-Gojobori method as implemented in MEGA 11. Percentile refers to rank relative to 10,766 genes compared between *D. melanogaster* and *D. simulans* (Stanley and Kulathinal 2016). Note that *mh* was not included in the Stanley and Kulathinal (2016) gene set because the *D. simulans* ortholog was not identified at that time.

### *D. simulans mh* complements *D. melanogaster mh* mutants

We constructed a transgene of *D. simulans mh* tagged with Gfp (called *sim-mh-Gfp*) to precisely match in structure the *D. melanogaster mh-Gfp* transgene of Tang et al. (2017). (Fig 1; Methods). We transformed and integrated both transgenes into *D. melanogaster* at the same autosomal position. Western blots indicate that both transgenes express at similar levels (Fig 2). We then crossed the transgenes into a *mh^6^* mutant background. Stable stocks were established, indicating that both transgenes can complement the sterility of *mh^6^*. To quantitatively compare the activity of these transgenes, we performed fertility assays of females trans-heterozygous for two *mh* null alleles and heterozygous for a *mh-Gfp* transgene; that is *w mh^6^ / w mh^31^; {mel-mh-Gfp, w^+^}/+* compared to *w mh^6^ / w mh^31^; {sim-mh-Gfp, w^+^}/+*. A small-scale pilot experiment found greater fertility among females carrying *mel-mh-Gfp*, but only among younger females (Fig 3A). We then performed a more extensive experiment, examining fertility across a 12-13 day period at two different temperatures (Fig 3B). No significant difference between the transgenic genotypes was observed at any time point, except for the first flip at 25°, where females carrying the *mel-mh-Gfp* transgene had significantly fewer progeny than those carrying the *sim-mh-Gfp* transgene. We conclude that the *D. melanogaster* and *D. simulans* orthologs have not substantially diverged for the essential female fertility function of *mh*.

**Fig 2.**
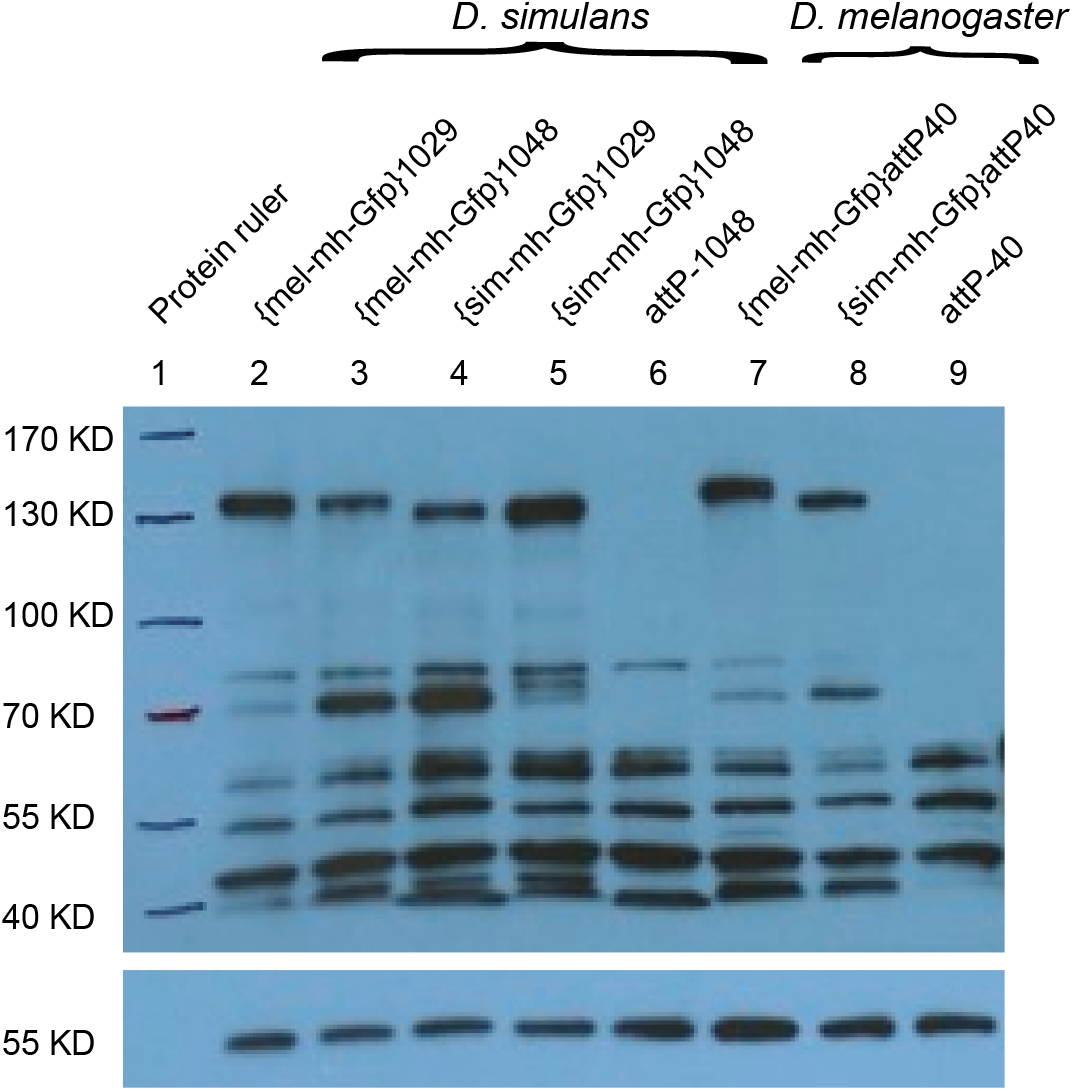
Western blot comparing mh-Gfp protein accumulation in different transgenic genotypes. Western blot with an anti-GFP antibody recognizes the Mh-eGFP protein in ovary extracts from females (top panel). Signals on the same membrane detected by an anti-α-Tubulin antibody served as a loading control (lower panel). Protein sizes of the pre-stained protein ladder were hand-marked on the membrane. Lanes 2-6 are from *D. simulans* extracts and lanes 7-9 from *D. melanogaster* extracts. Two independent transformants of the transgenes in *D. simulans* were analyzed, at *attP* sites 1029 and 1048. The *attP-1048* and *attP-40* samples are negative controls because they are the untransformed strains that carry the attP sites. The predicted molecular weights of mel-Mh-Gfp and sim-Mh-Gfp proteins are 108.6 and 109 kD, respectively.

**Fig. 3.**
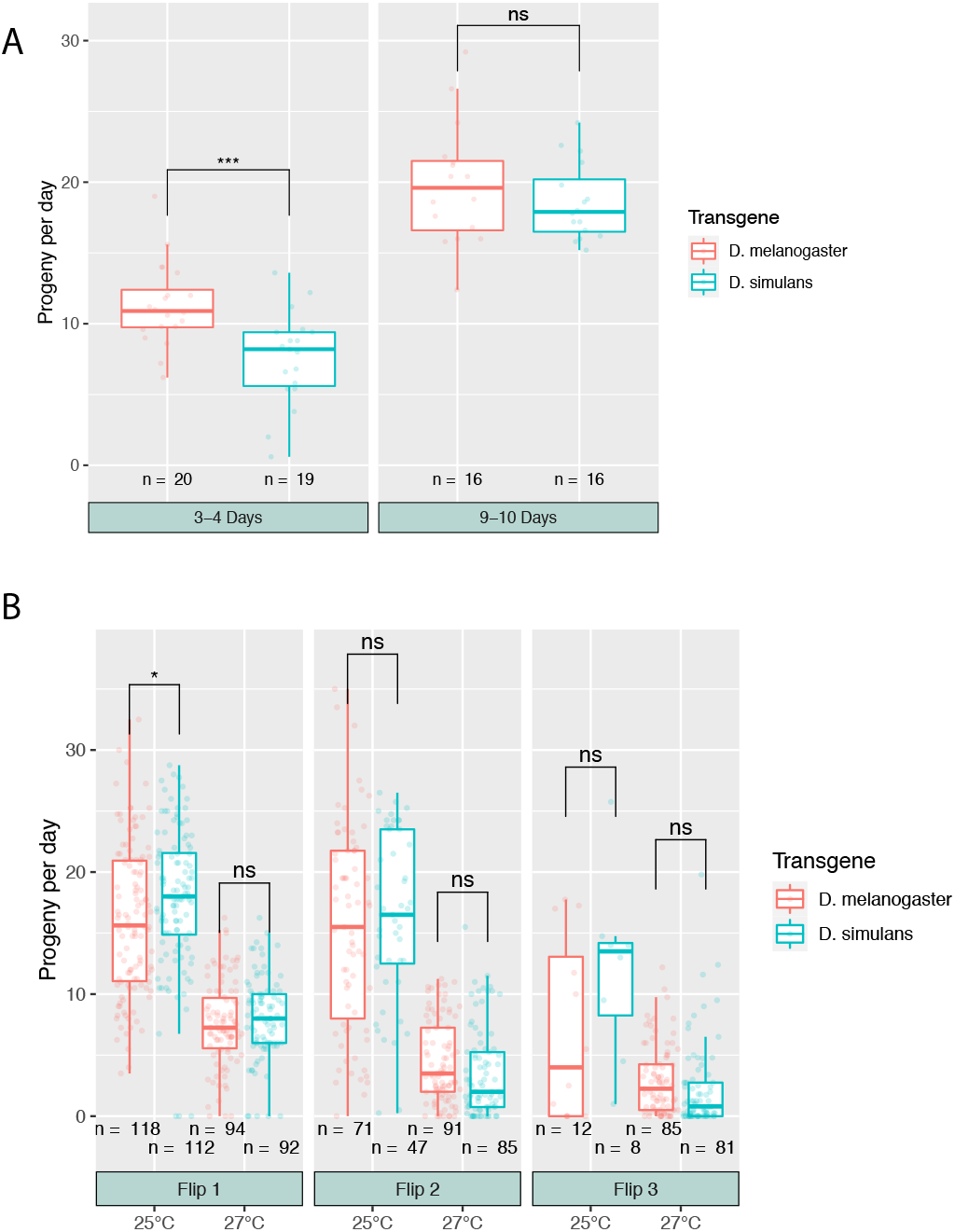
Comparison of fertility between mh mutant females carrying mel-mh-Gfp versus sim-mh-Gfp transgenes (single w mh6 / w mh31; {mel-mh-Gfp, w+}attP40/+ versus w mh6 / w mh31; {sim-mh-Gfp, w+}attP40/+). Statistical significance between genotypes was tested by a two-sample unpaired t-test: *** = 0.0001 < p ≤ 0.001; * = 0.01 < p ≤ 0.05; ns = p > 0.05 (not significant). A) Pilot experiment at room temperature (~20°-22°). Females were collected as virgins and aged for either 3-4 days (left) or 9-10 days (right) prior to mating and progeny collection for 5 days. Progeny are reported as ‘per-day’ to normalize with Fig 3B. B) Two large-scale experiments, performed at 25° and 27°. Virgin females were aged for 3-5 days prior to mating. Crosses were flipped to fresh vials after 4 or 5 days, and again after an additional 4 or 5 days, for a total collection period of 12-13 days; the exact length of each flip was recorded and is accounted for in calculating the ‘progeny per day’. Note that vials were discarded if the parents died, which is why N goes down between flips (see Methods).

### *D. melanogaster mh* does not affect hybrid viability

Having established that *sim-mh-Gfp* is functional within *D. melanogaster*, we turned to the primary question motivating this study, of whether *mh* is a hybrid incompatibility gene. We tested this by comparing the *D. simulans* and *D. melanogaster mh* transgenes for their ability to modulate F1 hybrid female viability. Specifically, we sought to determine whether the addition of a *mh^+^* transgene to the mothers of these hybrids would increase viability of their daughters. We transformed these transgenes into two different *attP* sites in *D. simulans*, with Western blots with an anti-Gfp antibody demonstrating similar protein levels for both transgenes (Fig 2).

We then generated F1 interspecific hybrids by crossing *D. simulans* females heterozygous for either *mel-mh-Gfp* or *sim-mh-Gfp*, along with sibling females not carrying a transgene as controls, to wild type *D. melanogaster* males (Table 2). The viability of the control F1 daughters not carrying a transgene was quite high, ranging from ~47%-69% relative to F1 sons, which limited our ability to detect substantial increases in the progeny of interspecific hybrids from mothers carrying a transgene. This relatively high rescue may reflect the fact that the transgenes were outcrossed to the *D. simulans w^501^* strain, which produced high hybrid viability in other crosses (Gérard and Presgraves 2012). Regardless, the results are opposite to our hypothesis. Both sets of crosses with the *sim-mh-Gfp* transgene showed increased female viability from transgenic mothers compared to control mothers, though only set D (with the 1048 insertion) was statistically significant. In contrast, crosses with the *mel-mh-Gfp* transgene showed essentially no differences in the relative viability of daughters between the transgenic and control genotypes. Within the resolution of our assay, we find no evidence suggesting that hybrid lethality results from the absence of *mel-mh* alleles in *D. simulans*.

**Table 2.** Testing *mh* transgenes for modulation of interspecific F1 hybrid viability

| Set | Maternal genotype | # F1 hybrid daughters | # F1 hybrid sons | Ratio F1 daughters/sons | Relative ratio w/o and w transgene | *P* value |
| --- | --- | --- | --- | --- | --- | --- |
| A | *w* | 760 | 1106 | 0.69 |  |  |
|  | *w; {mel-mh-Gfp}1029/+* | 766 | 1105 | 0.69 | 0.99 | 0.895 |
| B | *w* | 781 | 1144 | 0.68 |  |  |
|  | *w; {mel-mh-Gfp}1048/+* | 830 | 1124 | 0.74 | 0.92 | 0.228 |
| C | *w* | 144 | 211 | 0.68 |  |  |
|  | *w; {sim-mh-Gfp}1029/+* | 84 | 88 | 0.95 | 0.71 | 0.0722 |
| D | *w* | 248 | 529 | 0.47 |  |  |
|  | *w; {sim-mh-Gfp}1048/+* | 461 | 693 | 0.67 | 0.70 | 0.000331 |

In the following descriptions, all flies are *D. simulans* unless noted otherwise. The designation {mh-Gfp, w+} refers to one of the four transgenic genotypes used in this table. The flies described in the “Maternal genotype” column were generated as follows:
- 1) $w^{501}$ virgin females were crossed to $y\ w/Y; \{mh-Gfp, w+\}$ males. - 2) $w^{501}/y\ w; \{mh-Gfp, w+\}/+$ virgin daughters were crossed to $w^{501}/Y$ males. - 3) For each set, $y? w/w^{501}; \{mh-Gfp, w+\}/+$ and $y? w/w^{501}$ virgin daughters were separately collected.
The females from step 3) were then mated to *D. melanogaster* Canton-S males. The full genotypes of the transgenes are noted in the Materials and Methods. *P* values were calculated using a 2x2 Chi-squared test.

## Discussion

The *mh* gene is interesting based on its unusual mutant phenotype of producing gynogenetic haploid embryos. It also displays a relatively high rate of coding sequence evolution, and has experienced two duplications in Drosophila. Our major motivation in this study derived from the association of mel-Mh with the X-linked 359-bp satellite DNA block that is present in *D. melanogaster* but absent in *D. simulans*. We used transgenic constructs to compare the activity of *mh* orthologs from *D. melanogaster* and *D. simulans* in the background of both species. We designed the *sim-mh-Gfp* transgene to be parallel in structure to a *mel-mh-Gfp* transgene previously shown to be functional. Our *sim-mh-Gfp* transgene complements *D. melanogaster mh* mutations, as evidenced by the ability to maintain a fertile *mh^6^; sim-mh-Gfp* stock.

In the course of this study, Brand and Levine (2022) independently published a study of the evolution of *mh* between *D. melanogaster* and *D. simulans*. They compared *mh* ortholog function by replacing the endogenous *D. melanogaster mh* locus with the *D. simulans* ortholog. They found that females of this replacement line have reduced fertility compared to the *D. melanogaster mh* control line, and further found that this reduced fertility is dependent on an intact *Zhr^+^* locus, which contains a large block of the 359 bp satellite. Below we summarize our results and compare them to those of Brand and Levine where appropriate.

### Similar fertility function of *mel-mh* and *sim-mh*

We observed some reduction of fertility in *D. melanogaster mh* mutants carrying *sim-mh-Gfp* compared to *mel-mh-Gfp* controls in one of two small scale initial experiments (Fig 3A). However, follow-up experiments at much larger scale failed to find any reduction (with one out of the six comparisons showing a modest but significant increase in fertility of *sim-mh-Gfp* relative to *mel-mh-Gfp* females; Fig 3B). We conclude that *D. melanogaster* and *D. simulans mh* orthologs are largely interchangeable for female fertility.

This contrasts with the result of Brand and Levine, who report a more than two-fold mean reduction in fertility of *D. melanogaster* females carrying *sim-mh* compared to *mel-mh*. One possible explanation for these differences is that the fertility of *mh* genotypes are inherently variable due to environmental (or other) variation, which is plausible for any genotype that may be sub-fertile but not completely sterile. Another possibility may be differences in experimental design of the fertility assays. We assayed females individually while Brand and Levine analyzed fertility in broods of four females. If a female (and/or the males they were mating with) died during the course of our experiment we could exclude that vial, while in the Brand and Levine design individual deaths may not have been recorded and would reduce the brood size and thus presumably the progeny count. We saw very high rates of death in one of our experiments (at 25°) that varied by genotype: *mel-mh-Gfp* crosses dropped from 118 to 71 from flip 1 to 2, while for *sim-mh-Gfp* the drop was much greater, from 112 to 47.

There are also significant differences in experimental design of the gene replacements between the two studies. Here we have used transgenic constructs integrated into an autosomal site and crossed into *mh* null-allele backgrounds to ‘replace’ *mh* and compare *mel-mh* and *sim-mh*. We designed our *sim-mh-Gfp* transgene to match a previously described *mel-mh-Gfp* transgene that was shown to provide wild type *mh* function (Tang *et al*. 2017). This introduces a potential position effect as *mh* is now in an autosomal location. It is also possible that *sim-mh-Gfp* does not express properly in a *D. melanogaster* background since it has its endogenous regulatory regions. We did, however, see robust protein expression from our transgenes. Brand and Levine (2022) used a CRISPR replacement strategy, replacing the endogenous *mh* locus in *D. melanogaster* with FLAG-tagged and codon-optimized *mel-mh* and *sim-mh* coding sequences. This means that the *sim-mh* allele is synthetic in the sense that the synonymous sites in the transgene are not native to *D. simulans*. But again, Western blots suggest that it is fully expressed. Among other differences in the two studies is that our fertility assays were done with one functional copy of *mh* in the mothers, while the Brand and Levine (2022) design has two functional copies. Brand and Levine (2022) found that *D. melanogaster* females with one *mel-mh* and one *sim-mh* allele are fully fertile, and other experiments support their conclusion that deleterious effects of *sim-mh* on fertility and ovarian morphology are dose-dependent.

### Lack of effect of *mel-mh* on hybrid viability

We found no evidence that the absence of *mel-mh* contributes to hybrid female lethality. In crosses of *D. simulans* females to *D. melanogaster* males, *D. simulans* mothers carrying a *mel-mh-Gfp* transgene produced the same ratio of female hybrids compared to control crosses without the transgene. It remains possible that repeating our experiments in *D. simulans* genetic backgrounds that have a lower baseline of hybrid female viability might reveal more subtle effects that we could not detect. Surprisingly, we did observe a moderate effect of increased hybrid female viability produced by mothers that carried the *sim-mh-Gfp* transgene. This finding suggests that increased *mh* dosage may reduce mis-segregation of the 359-bp satellite in hybrids, but that such effects may not be dependent on the *mel-mh* ortholog that has co-evolved with the *D. melanogaster* 359-bp satellite block.

## Acknowledgements

Plasmids were obtained from the Drosophila Genomics Resource Center (supported by NIH Grant 2P40OD010949). The w+attB plasmid was a gift from Jeff Sekelsky (Addgene plasmid # 30326). We thank David Stern (HHMI Janelia Farms) for sharing *D. simulans attP* strains. Supported by NIH R01-GM07473 to DAB. DMC was supported by an NIH NRSA fellowship 5F32GM120896.

---

**Table S1.**
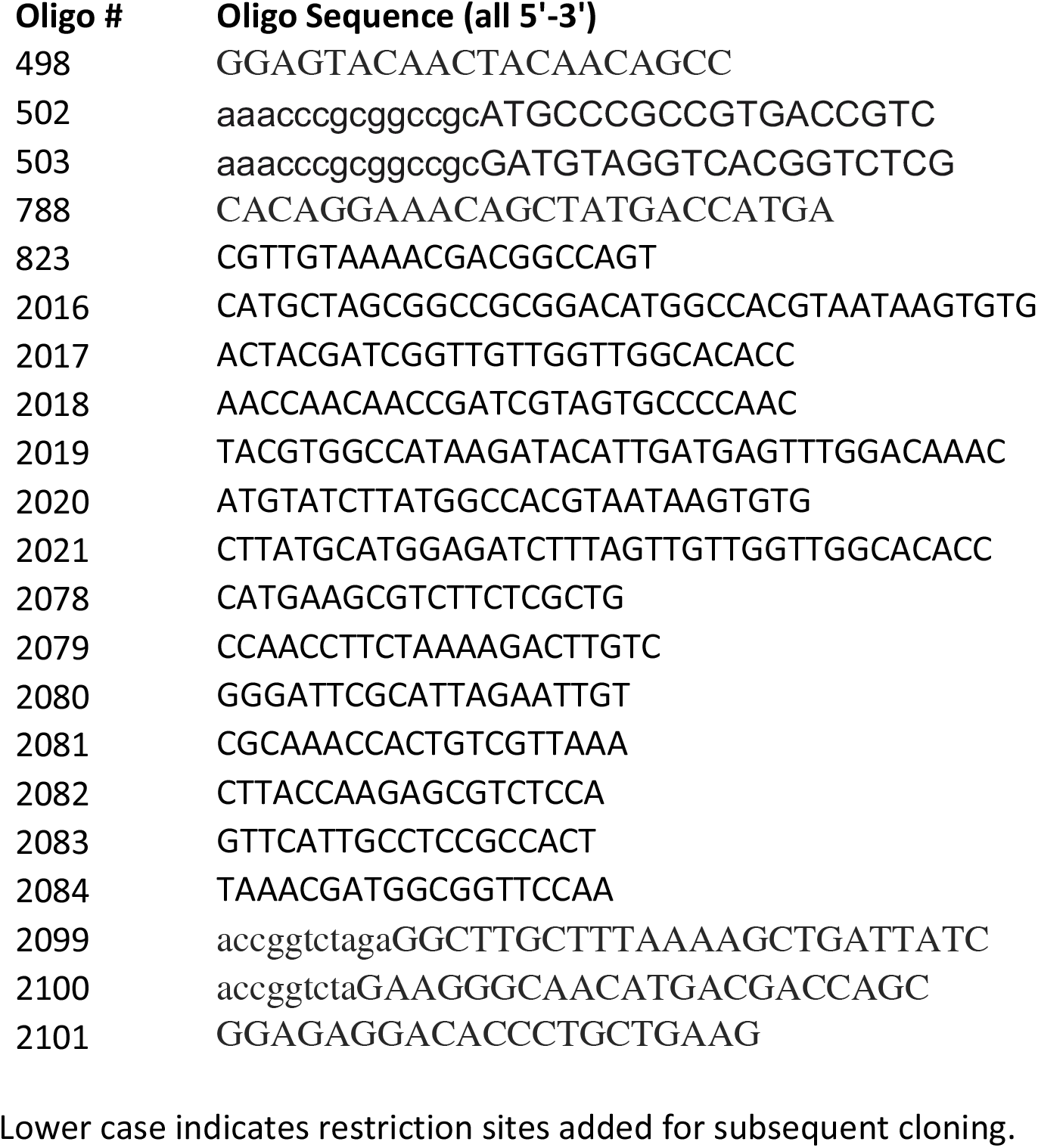
Oligonucleotide sequences

